# Activity-induced Ca^2+^ signaling in perisynaptic Schwann cells is mediated by P2Y_1_ receptors and regulates muscle fatigue

**DOI:** 10.1101/176354

**Authors:** Dante J. Heredia, Cheng-Yuan Feng, Grant W. Hennig, Thomas W. Gould

## Abstract

Perisynaptic glial cells respond to neural activity by increasing cytosolic levels of calcium, but the functional significance of this pathway is unclear. Terminal/persiynaptic Schwann cells (TPSCs) are a perisynaptic glial cell at the neuromuscular junction. Here, we provide genetic evidence that neural activity-induced intracellular calcium accumulation in neonatal TPSCs is mediated exclusively by P2Y_1_ receptors. In *P2ry1* mutant mice lacking these responses, postsynaptic, rather than presynaptic, function was altered in response to nerve stimulation. This impairment was correlated with a greater susceptibility to activity-induced muscle fatigue. Interestingly, fatigue in *P2ry1* mutants was exacerbated by exposure to high potassium to a greater degree than in control mice. High potassium itself increased cytosolic levels of calcium in TPSCs, a response which was also reduced *P2ry1* mutants. These results suggest that activity-induced calcium responses in perisynaptic glia at the NMJ regulate postsynaptic function and muscle fatigue by influencing the levels of perisynaptic potassium.

## Introduction

Muscle fatigue is defined as the decline in muscle performance that occurs in response to continued muscle activation. Muscle fatigue is a clinically important feature of myopathies such as muscular dystrophy, neuromuscular disorders such as Guillain-Barré syndrome, diseases of the central nervous system (CNS) such as multiple sclerosis, or diffuse conditions such as chronic fatigue syndrome and cachexia (Katirji, 2002). A wide variety of mechanisms in the central and peripheral nervous systems contribute to muscle fatigue. In simplified preparations of muscle and peripheral nerve, central sources of input are eliminated, permitting the examination of peripheral sites of fatigue, such as the presynaptic release of the neurotransmitter acetylcholine (ACh) at the neuromuscular junction (NMJ; Nanou et a., 2016), sensitivity of postsynaptic ACh receptors, propagation of the muscle action potential along the sarcolemma and into the t-tubules, release of calcium (Ca^^2^+^) from sarcoplasmic reticulum, and activation of the contractile apparatus (Boyas and Guével, 2011). Proposed mediators of fatigue at these sites include changes in the concentration of intracellular and extracellular ions, such as calcium (Ca^^2^+^), sodium (Na^+^), potassium (K^+^), or protons (H^+^); metabolites, such as inorganic phosphate (P_i_), or lactate; and reactive oxygen species (Allen et al., 2008). For example, high-frequency stimulation (HFS) of nerve or muscle raises the level of extracellular K^+^ or [K^+^]_o_, which may mediate fatigue by depolarizing muscle membrane, inactivating Na_v_1.4 voltage-gated sodium channels at the NMJ, and consequently blocking the production of muscle action potentials (APs; Cairns et al., 2015) This mechanism may also underlie the muscle weakness observed in patients with hyperkalemic periodic paralysis, a neuromuscular disorder caused by dominant mutations in the *Scna4* gene encoding the Na_v_1.4 channel and characterized by episodic muscle stiffness and weakness (Cannon, 2015).

In addition to presynaptic nerve terminals and muscle endplates, terminal or perisynaptic Schwann cells (TPSCs) reside at the NMJ. TPSCs are a non-myelinating Schwann cell subtype that influence the regeneration of injured peripheral motor axons (Son et al., 1996), maintain developing synapses (Reddy et al., 2003), and participate in synaptic pruning (Smith et al., 2013). TPSCs also respond to neural activity by increasing cytosolic Ca^^2^+^ levels (Jahromi et al., 1992; Reist and Smith, 1992) and are therefore functionally similar to other perisynaptic glial cells, such as astrocytes and enteric glia. In addition to responding to neurotransmitter released during neural activity by mobilizing Ca^^2^+^, astrocytes regulate the concentration of extracellular metabolites produced by activity through the expression of various ion channels and transporters (Olsen et al., 2015; Boscia et al., 2016; Weller et al., 2016). Therefore, TPSCs, as the perisynaptic glia of the NMJ, likely act to modulate the concentrations of these ions at the NMJ and thereby regulate muscle fatigue.

In astrocytes, activity-induced Ca^^2^+^ signaling is largely mediated by neurotransmitter-mediated stimulation of Gq G-protein coupled receptors (GPCRs), leading to the release of Ca^^2^+^ from the endoplasmic reticulum (ER) through the second messenger inositol-1,4,5-triphosphate (IP_3_; Volterra et al., 2014). Astrocytic Ca^^2^+^ signaling in turn modulates synaptic transmission and contributes to functional hyperemia, although each of these effects remains controversial (Agulhon et al., 2010; Bonder and McCarthy, 2014). The interpretation of these effects is complicated by the observation that the mechanisms contributing to activity-induced Ca^^2^+^ signaling in the fine processes of astrocytes are distinct from those underlying this signal in the cell body (Bazargani and Atwell, 2016). Additionally, the diversity of astrocyte subtypes in the brain (Eugenín León et al., 2016) and of neuronal subtypes associated with individual astrocytes (Perea and Araque, 2005) further challenge the precise identification of activity-induced Ca^^2^+^ responses in these cells. Finally, the extent to which IP_3_R-mediated Ca^^2^+^ signaling reflects all of the effects of activity-induced Gq GPCR activation remain unclear (Agulhon et al., 2013).

TPSCs, by contrast, are only associated with the nerve terminals of cholinergic motor neurons (MNs). The NMJ is large and amenable to optical analysis owing to its discrete location at the central endplate band region of muscle. TPSCs do not elaborate extensive processes or make specialized contacts with the microcirculation. Together, these features make the TPSC suitable for the genetic manipulation of their response to and regulation of neural activity. However, the examination of activity-induced Ca^^2^+^ responses in TPSCs and the functional effects of these responses have been largely conducted by imaging individual TPSCs injected with fluorescent Ca^^2^+^-binding dyes (Darabid et al., 2014). Although this approach allows for the evaluation of TPSC function at individual synapses, the extent to which these mechanisms are shared by all TPSCs is unclear. Moreover, whether the effects of single TPSC manipulation on individual synapses lead to global effects on neuromuscular function cannot be examined.

Using mice that express the genetically-encoded calcium indicator GCaMP3 in all Schwann cells including TPSCs, we observed that high-frequency, motor nerve stimulation-induced Ca^^2^+^ signaling within TPSCs of the neonatal diaphragm was completely abolished in the absence of the purinergic 2Y_1_ receptor (P2Y_1_R). We therefore utilized *P2ry1* mutant mice lacking these receptors as a model to investigate the functional effects of activity-induced, Gq GPCR-mediated Ca^^2^+^ release in TPSCs.

## Results

### Activity-induced Ca^2+^ transients in populations of TPSCs in Wnt1-GCaMP3 mice

In order to study the Ca^2+^ response to neural activity in populations of TPSCs, we first evaluated transgene expression in TPSCs of the diaphragm muscle at postnatal day 7 (P7) from Wnt1-Cre, conditional GCaMP3 (Wnt1-GCaMP3) mice. Wnt1-Cre mice drive Cre-dependent transgene expression in neural crest derivatives, which include Schwann cells (Danielian et al., 1998). We first assessed Wnt1-Cre mice by crossing them to mice conditionally expressing the fluorescent reporter TdTomato. Robust expression of TdTomato was observed at early ages in all Schwann cells in the diaphragm, including myelinating Schwann cells of the phrenic nerve as well as non-myelinating TPSCs at the NMJ (**Figure 1A**), visualized using fluorescent α–bungarotoxin (α-BTX). We then crossed Wnt1-Cre mice to mice conditionally expressing GCaMP3, which encodes a green fluorescent protein (GFP)-conjugated calmodulin that fluoresces upon Ca^2+^ binding (Zariwala et al., 2012). In whole-mounts of P7 diaphragm muscle incubated with GFP antibodies to label GCaMP3, S100 antibodies to detect Schwann cells, and α-BTX to visualize NMJs, we observed expression of GCaMP3 in all TPSCs (**Figure 1B**). Together, these results show that Wnt1-Cre drives robust expression of GCaMP3 in TPSCs of the early postnatal diaphragm.

**Figure 1.**
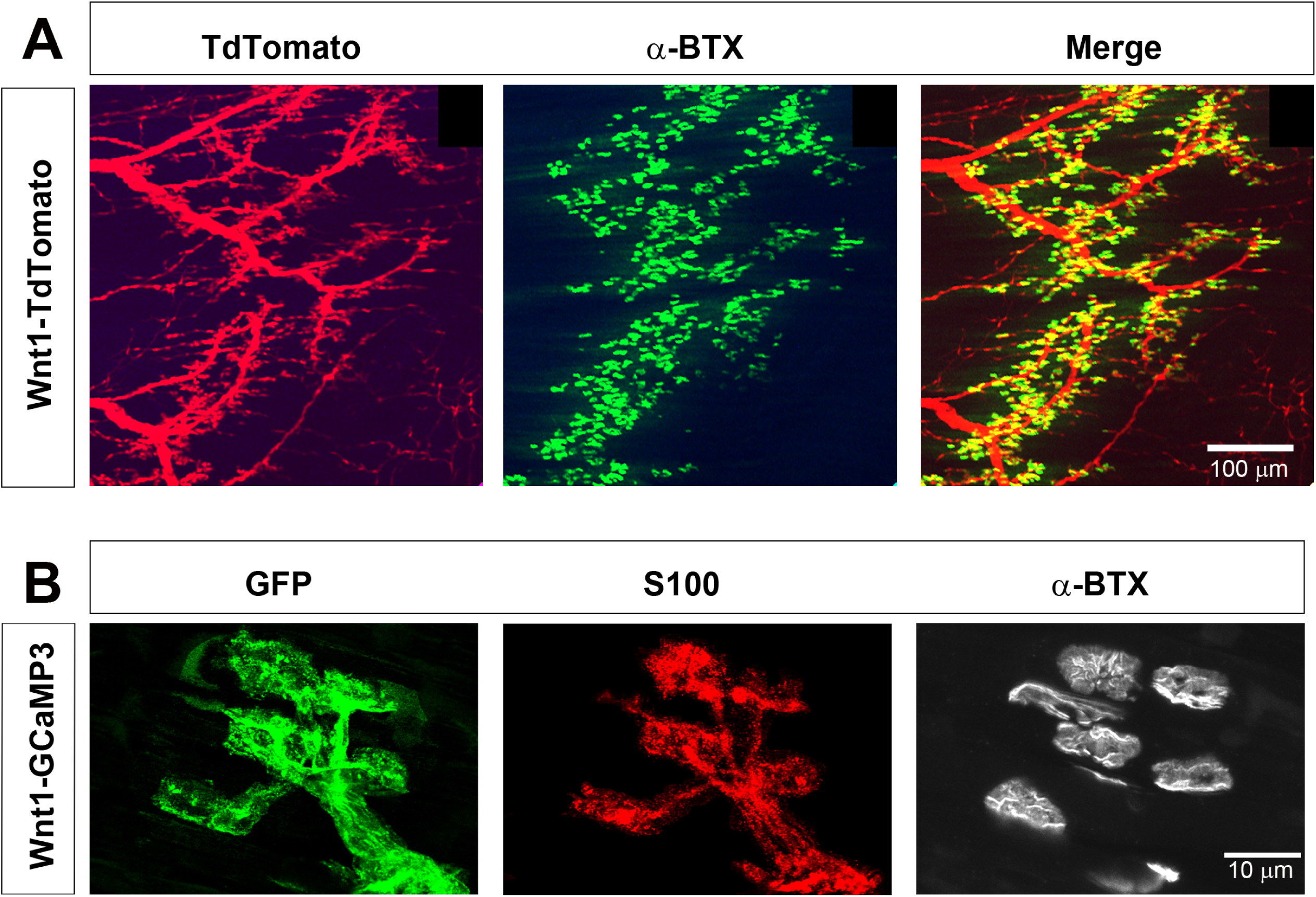
Wnt1-Cre drives expression of reporters and activity sensors to Schwann cells of the neonatal diaphragm. ***A***, Whole mounts of P7 Wnt1-TdTomato diaphragm were labeled with 488-conjugated α-bungarotoxin (α-BTX). ***B***, Higher magnification of whole mounts of P7 Wnt1-GCaMP3 diaphragm labeled with GFP, S100, and 633-conjugated α-BTX. All NMJ-associated, S100-immunostained TPSCs express GFP and thus GCaMP3.

We next determined if GCaMP3 expression in TPSCs exhibited activity-induced Ca^2+^ responses, similar to previous studies (Jahromi et al., 1992; Reist and Smith, 1992). Imaging these responses before and after nerve stimulation at low magnification (20X), we observed large populations of TPSCs that responded to 45s of 40Hz tonic phrenic nerve stimulation (**Supplemental Video 1; Figure 2A**). Higher magnification images (60X) showed that each individual TPSC, identified by labeling with fluorescent α-BTX (data not shown), responded to HFS (**Figure 2B**). We used stat maps of the standard deviation of fluorescence intensity (SD map) to spatially represent the distribution of Ca^2+^ transients within individual TPSCs from high-magnification videos and traces of intensity to examine their temporal characteristics (**Figure 2C**). The onset of these transients occurred 1.8 ± 0.74s (*n*=5, *c*= 54) after the initiation of HFS, reached peak intensity at 2.86 ± 0.49s after the onset of the transient, and shortly thereafter declined in amplitude (time to 50% decay = 13.35 ± 4.22s). Most transients lasted the entire duration of the nerve stimulation period, although the intensity at the end of the stimulation period was usually less than 10% of that at the initial peak. GCaMP3-expressing TPSCs responded to multiple stimulations, providing a useful tool to measure the effects of different stimuli on the same TPSCs. Similar to a previous study (Darabid et al., 2013), the peak intensity of Ca^2+^ transients observed after a subsequent 45s bout of 40Hz stimulation was lower than that of the first (33 ± 4.1 vs. 26.5 ± 6.1 dB, first vs. second stim, *n*=5; *p*<0.01). Interestingly, analyses of transients in TPSCs across large regions of the diaphragm from low-magnification videos showed substantial variability in the onset after nerve stimulation (from 0.8s to 4.3s). These results are not related to heterogeneity in the onset of neurotransmitter release, as individual muscle fibers across the entire diaphragm exhibited shortening at nearly the same onset after nerve stimulation (**Supplemental Video 1**).

**Figure 2.**
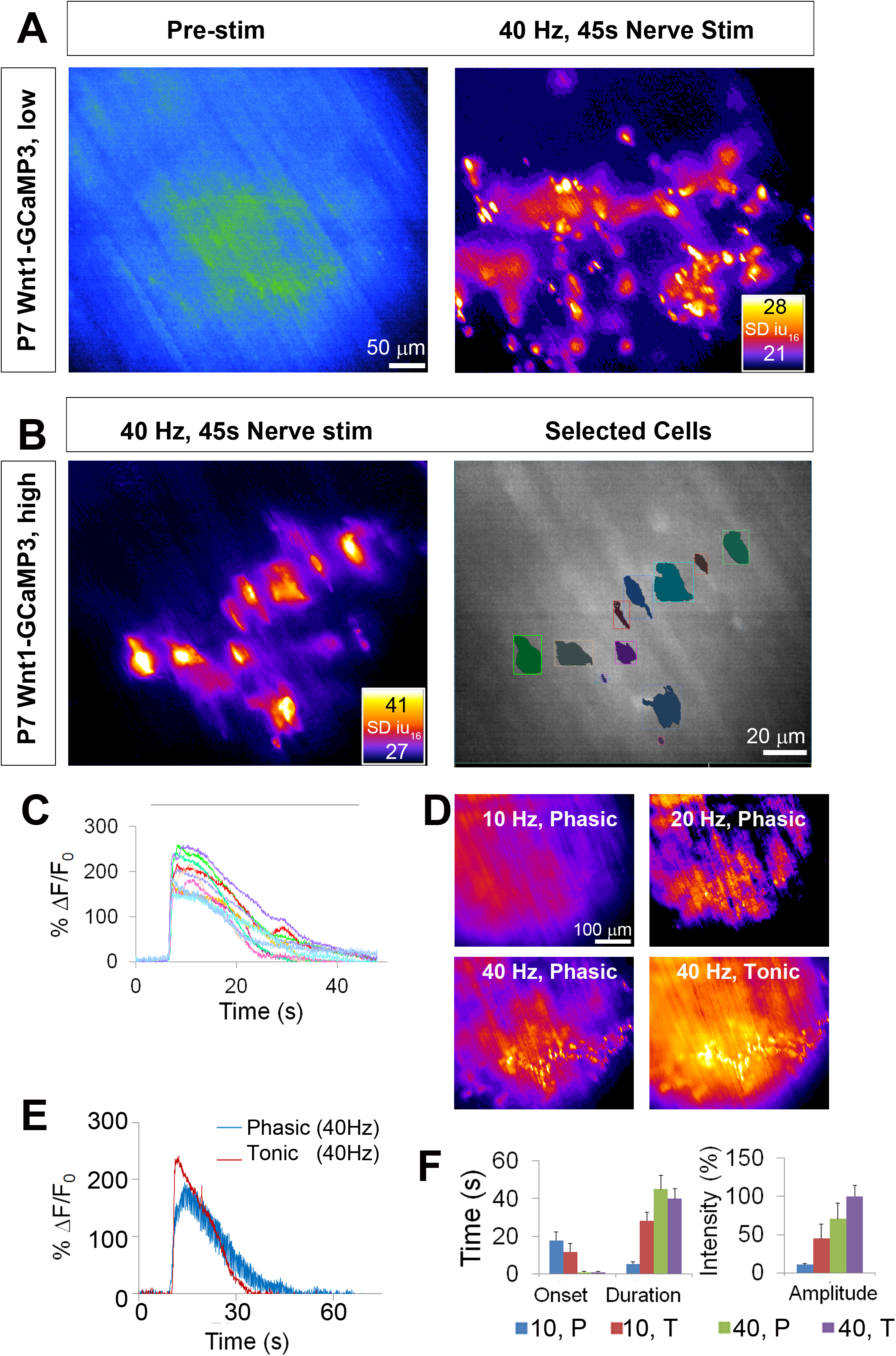
Wnt1-GCaMP3 mice exhibit activity-induced Ca^2+^ responses in all perisynaptic glia cells of the neonatal diaphragm. ***A***, (Left panel) An average intensity image generated before application of a stimulus (Pre-stim) shows the overall structure of GCaMP3-expressing Schwann cell elements. (Right panel) Map of standard deviation of 16-bit fluorescence intensity units (SD iu_16_) of a population of TPSCs imaged in response to high-frequency nerve stimulation at low magnification; fire CLUT heatmap in SD iu_16_. ***B***, Same muscle imaged at higher power showing these fluorescence responses in individual TPSCs (left panel) were color coded (right panel). ***C***, These cells in B were plotted as color-coded transients. ***D***, SD maps resulting from tonic vs. phasic HFS. ***E***, Transients elicited by 40 Hz tonic (red) or (blue) phasic stimulation. Peak transient intensity was lower (22.9 ± 3.2 vs. 16.2 ± 5.1 dB; *p*<0.05; *n*=3; *c*=18), and duration was longer (time to 50% decay = 19.4 ± 4.6 vs. 14.7 ± 6.2s; *p*<0.05) in response to phasic vs. tonic stimulation. ***F***, Slower onsets and lower amplitudes of Ca^2+^ transients in TPSCs stimulated with 10 vs. 40 Hz stimulation (10Hz phasic = 17.92 ± 6.3; 10Hz tonic = 11.78 ± 3.4s; 40Hz phasic = 12.1 ± 0.91s; 40Hz tonic = 1.8 ± 0.74s). 10=10Hz; 40=40Hz; T=tonic; P=phasic.

Although the diaphragm is activated by short bursts of tonic stimulation of the phrenic nerve during several behaviors (e.g., expulsive maneuvers such as wretching; Hodges and Gandevia, 2000), the more native physiological pattern of stimulation that occurs during respiration is phasic with a duty cycle between 25 and 50%, in which each period of activation is 100ms, at frequencies between 30-70Hz (Kong and Berger, 1986; Zhan et al., 1998; van Lunteren and Moyer, 2003; Sieck et al., 2012). The peak intensity of TPSC Ca^2+^ transients was lower and the duration was longer in response to 45s of 40 Hz phasic vs. tonic stimulation (**Figure 2E,F**). The off-period of each duty cycle could clearly be discerned as a dropoff in transient intensity (**Figure 2E**), showing the dynamic nature of these responses (Todd et al., 2010; Darabid et al., 2013). At lower frequencies of stimulation, we observed similar differences of Ca^2+^ signals in response to phasic vs. tonic patterns. Interestingly, in response to 10Hz stimulation, the lowest phasic rate that produced a measurable response, Ca^2+^ transients were much slower in onset after nerve stimulation than after 40Hz stimulation (**Figure 2F**).

We next examined whether lower frequencies were capable of inducing Ca^2+^ transients in TPSCs, similar to astrocytes (Sun et al., 2014). Whereas spontaneous ACh release, 1Hz and 2Hz evoked stimulation failed to produce visible Ca^2+^ transients in TPSCs, 5Hz stimulation elicited transients in a small number of TPSCs, and 10-40Hz stimulation produced responses in all TPSCs (data not shown). In response to different durations of 40Hz stimulation, only several TPSCs responded to 0.1s of nerve stimulation (i.e., 4 pulses), whereas all TPSCs responded to 1-30s of 40Hz stimulation (**Supplemental Video 2**). Finally, in contrast to TPSCs, myelinating Schwann cells along phrenic nerve trunks and branches failed to exhibit Ca^2+^ responses, similar to previous reports (Jahromi et al., 1992). However, bath application of adenosine triphosphate (ATP) or muscarine elicited a response in Schwann cells along distal phrenic nerve branches as well as TPSCs (**Supplemental Videos 3,4**). Despite their expression of GCaMP3 (determined by immunohistochemistry; data not shown), myelinating Schwann cells along larger phrenic nerve branches failed to respond to either nerve stimulation or bath application of ATP or muscarine. Together, these results show that TPSCs dynamically respond to different patterns of neuronal activity.

### Activity-induced Ca^2+^ responses in TPSCs of the neonatal diaphragm are mediated exclusively by purinergic stimulation of P2Y_1_ receptors

Previous studies of individual or small cohorts of TPSCs show that a variety of neurotransmitter-derived substances trigger cytosolic Ca^2+^ accumulation, including ACh through muscarinic AChRs, adenine nucleotides such as ATP/ADP through purinergic P2Y receptors (P2YR), and adenosine, derived from synaptic ectonucleotidase-mediated degradation of adenine nucleotides (Cunha et al., 1996), through P1R (Darabid et al., 2014). A recent study provided evidence that TPSCs also respond to nerve-derived ACh through nicotinic AChRs (Petrov et al., 2014). In contrast to these responses in adult TPSCs, a recent report demonstarted that TPSC Ca^2+^ signals in neonatal mouse soleus are not mediated by muscarinic or nicotinic ACh receptor (mAChR) or by adenosine receptor activation, but rather by P2YRs (Darabid et al., 2013). In agreement with this finding, we found that whereas the pan-muscarinic antagonist atropine and the pan-nicotinic antagonist curare failed to block activity-mediated Ca^2+^ transients in TPSCs of the P7 diaphragm (*n*=8), the pan-P2 antagonist suramin completely eliminated them (*n*=6; **Figure 3C**). We tested whether this response was mediated by P2Y_1_ receptors (P2Y_1_Rs), as TPSCs reportedly express this protein (Darabid et al., 2013) and astrocytic Ca^2+^ signaling is mediated in part by the Gq GPCR-coupled pathway that is activated by P2Y_1_Rs (Fam et al., 2002). Treatment with the selective P2Y_1_R antagonist MRS2500 (1μM) completely blocked activity-induced Ca^2+^ responses in all TPSCs of the diaphragm (*n*=5; **Figure 3C**). In order to determine whether activity-dependent, P2Y_1_R-mediated Ca^2+^ responses were dependent on release from intracellular stores, we examined them in the presence of the sarco-/endoplasmic reticulum Ca^2+-^ATPase (SERCA) inhibitor cyclopiazonic acid (CPA). 15 minutes after treatment with CPA, these responses were completely abolished (*n*=3; **Figure 3C**). Finally, to test whether the effect of MRS2500 was indeed mediated by P2Y_1_Rs, we crossed mice expressing a constitutive null mutation of the *P2ry1* gene, which encodes P2Y_1_Rs (Fabre et al., 1999), to Wnt1-GCaMP3 mice. Nerve stimulation of *P2ry1* mutants completely failed to elicit Ca^2+^ responses in TPSCs (*n*=7; **Supplemental Video 4, Figure 3A,C**). Mutant TPSCs exhibited a robust response to muscarine (**Figure 3A,C**), indicating that the failure of activity to induce these responses was not caused by non-specific effects of the *P2ry1* mutation.

**Figure 3.**
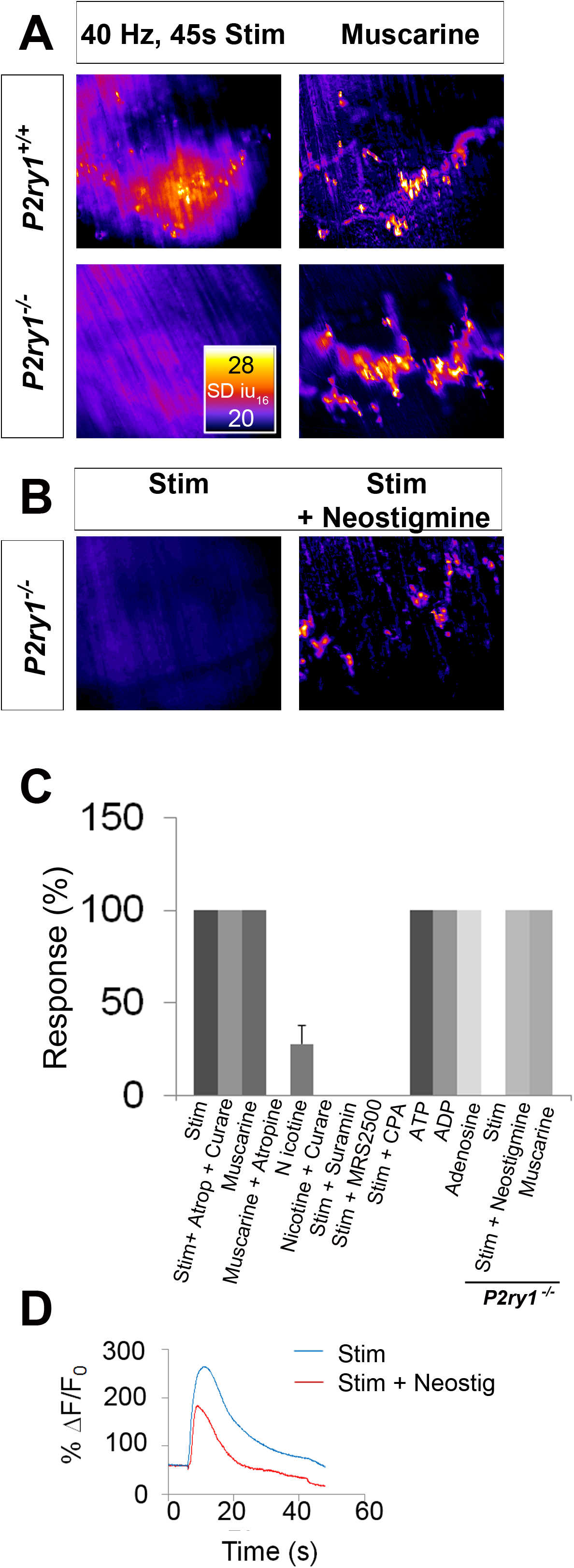
Activity-induced responses in perisynaptic glia at the NMJ are completely dependent on P2Y_1_R signaling. ***A***, SD maps of activity-induced (left panels) or muscarine-induced (right panels) Ca^2+^ responses in TPSCs of *P2ry1* wild-type (WT; top panels) or mutant (bottom panels) diaphragm. Note the complete absence of activity-induced Ca^2+^ responses in TPSCs of *P2ry1* mutants; fire CLUT heatmap in SD iu_16_. ***B***, Addition of neostigmine restores this response. ***C***, Graph representing the relative number of TPSCs exhibiting Ca^2+^ responses in response to activity or drug treatment. ***D***, WT mice exhibit elevated Ca^2+^ responses to activity in the presence of the cholinesterase-blocking drug neostigmine (33.3 ± 4.2 vs. 29.1 ± 6.6 dB; *p*<0.05; *n*=3; *c*=22).

We were intrigued by the inability of atropine to block activity-mediated Ca^2+^ responses in TPSCs of *P2ry1* WT mice, since a.) bath application of muscarine evoked a robust Ca^2+^ signal.; b.) atropine blocked this effect of bath-applied muscarine (data not shown); c.) ACh is released upon nerve stimulation. On the one hand, this may result from the absence of clustered mAChRs in P7 TPSCs (Darabid et al., 2013). Alternatively, the lateral diffusion of nerve-derived ACh to perisynaptic TPSC-derived mAChRs may be limited by the activity of the cholinesterases acetylcholinesterase (AChE) or butyrylcholinesterase (BChE) in the synaptic cleft. Nerve stimulation of *P2ry1* mutants in the presence of the pan-cholinesterase inhibitor neostigmine resulted in a robust Ca^2+^ response in all TPSCs, suggesting that cholinesterase activity normally prevents this effect (*n*=3; **Figure 3B,C**). Neostigmine also increased the intensity of activity-induced Ca^2+^ transients in *P2ry1* WT mice, demonstrating the additive nature of the effects of purinergic and muscarinic stimulation on this response (**Figure 3D**).

We investigated which nerve-derived, P2Y_1_R-stimulating ligands were capable of evoking TPSC Ca^2+^ responses. While bath application of either ATP or ADP induced these responses, neither ADP-ribose nor β-nicotinamide adenine dinucleotide (βNAD) did (*n*=4; data not shown; Mutafova-Yambolieva and Durnin, 2014). Bath application of the P1R-activating ligand adenosine also evoked robust Ca^2+^ transients in TPSCs (*n*=3; data not shown), similar to previous studies (Robitaille, 1995; Castonguay and Robitaille, 2002). However, the onset of this response was markedly delayed, compared to that triggered by purines or ACh mimetics (ATP = 3.2 ± 1.6s, adenosine = 15.6 ± 3.4s; *p*<0.001). Together, these pharmacological and genetic studies suggest that activity-induced Ca^2+^ signaling in neonatal diaphragm TPSCs is exclusively mediated by nerve-derived, adenine nucleotide-mediated stimulation of P2Y_1_R. This result thus permits the evaluation of the functional role of this signal.

### Activity-induced, P2Y_1_R-mediated Ca^2+^ responses in TPSCs are not required for synapse formation

In order to test whether P2Y_1_R deletion itself or activity-induced P2Y_1_R-mediated Ca^2+^ signaling exerted gross effects on synaptogenesis of the NMJ, we first assessed the structure of NMJs by immunohistochemical and ultrastructural techniques. We examined the tripartite NMJ by staining whole-mounts of P7 diaphragm with antibodies against synaptophysin (nerve terminals), GFP (GCaMP3-expressing perisynaptic Schwann cells) and α-BTX (postsynaptic AChR clusters). We found no difference in the total number of NMJs, the size of NMJs, the percentage of innervated NMJs, or the apposition of perisynaptic Schwann cells in *P2ry1* mutant or WT mice (**Supplemental Figure 1A,B**; data not shown). Next, we examined NMJs for AChE immunoreactivity, as previous reports of adult NMJs in *P2ry1* mutants showed a reduction in the level of expression of this cholinesterase (Xu et al., 2015). Using a highly specific antibody that fails to detect expression of this enzyme in *AChE* mutant mice, we were unable to observe any difference in its expression or synaptic localization (**Supplemental Figure 1C**). Finally, we examined NMJs by electron microscopy, and found no obvious structural abnormalities (**Supplemental Figure 2**). Together, these results suggest that the gross morphological development of the NMJ, at least until P7 in the diaphragm, is unaffected in mice lacking P2Y_1_R and activity-induced Ca^2+^ signaling.

### Activity-induced, P2Y_1_R-mediated Ca^2+^ responses in TPSCs are required for postsynaptic function in response to HFS

We next evaluated the effect of eliminating activity-induced Ca^2+^ signaling on the presynaptic release of neurotransmitter, based on results obtained in previous studies (Robitaille, 1998; Castonguay and Robitaille, 2001). The resting membrane potential (RMP) was unchanged between genotypes (-68.5 ± 5.2 vs. -68.4 ± 3.3mV; WT vs. mutant; p=0.46; WT vs. mutant; n=3, c=13), and was not significantly different after vs. before a nerve stimulation bout between genotypes (data not shown). The frequency, amplitude, rise to peak, and time to 50% decay of miniature EPPs (mEPPs) were also unchanged (resting frequency: 0.38 ± 0.15 vs. 0.46 ± 0.14 events/s; p=0.11; post-stimulation frequency: 1.57 ± 0.8 vs. 2.01 ± 0.8 events/s; p=0.15; amplitude: 2.52 ± 0.5 vs. 2.49 ± 0.6mV; p=0.44; rise to peak: 3.76 ± 1.8 vs. 3.69 ± 2.7ms; p=0.46; time to 50% decay: 5.83 ± 2.4 vs. 4.81 ± 1.8ms; p=0.08; WT vs. mutant; *n*=3, *c*=13-19). Individual nerve-evoked EPPs, recorded in the presence of μ-conotoxin, were also similar between *P2ry1* WT and mutant mice (**Figure 4A**). In response to HFS, EPP amplitudes at the end of the period were also similar in each genotype. These results demonstrate that basal and HFS-induced ACh release are not affected in the absence of activity-induced Ca^2+^ signaling in TPSCs (**Figure 4A,D**). These results also corroborate the finding that AChE expression at the NMJ was unaffected in *P2ry1* mutants, as the durations of mEPPs and EPPs were unaffected, whereas they are longer in the absence of this enzyme (Adler et al., 2011).

**Figure 4.**
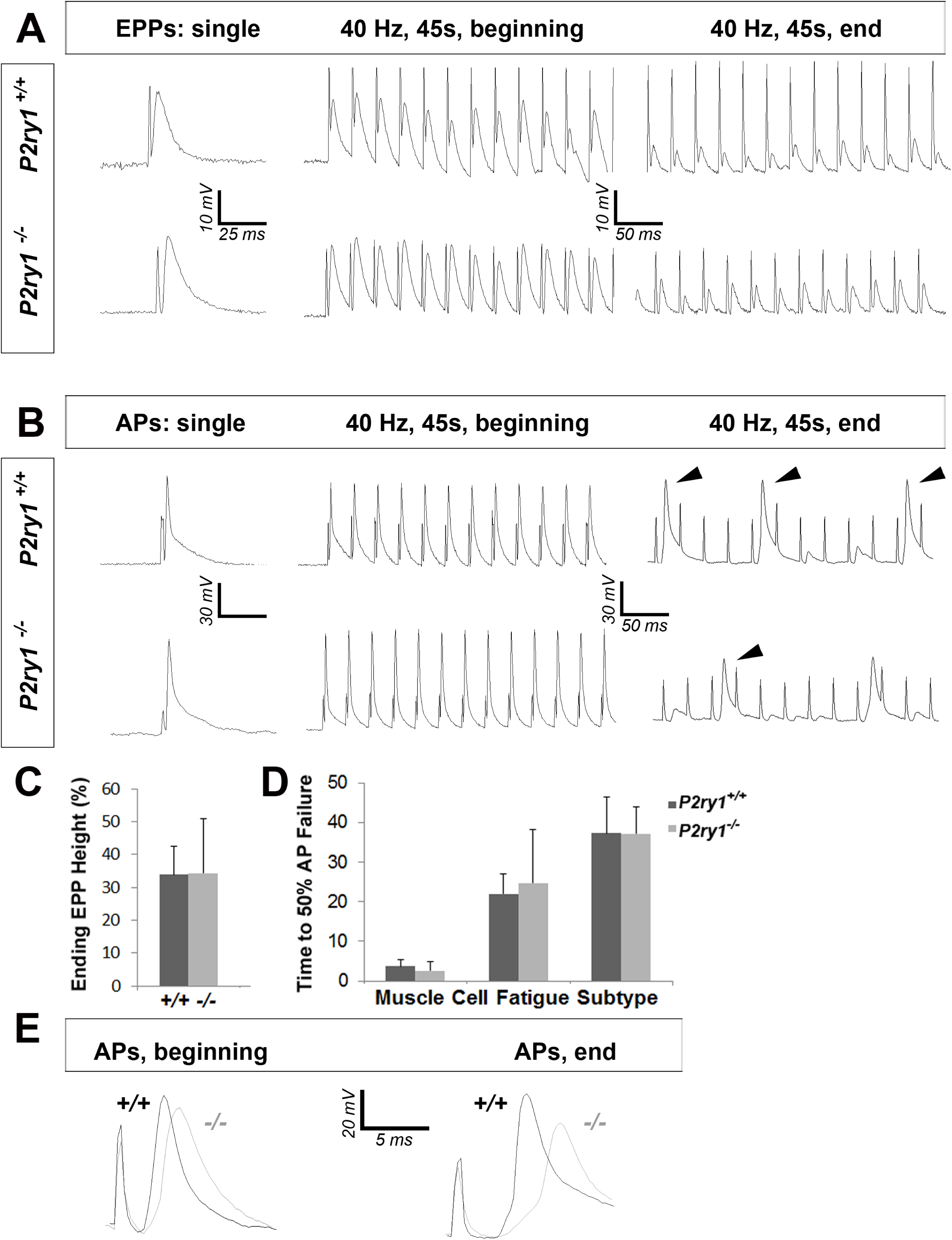
The loss of activity-induced Ca^2+^ responses in perisynaptic glia disrupts postsynaptic but not presynaptic function at the NMJ. ***A***, Phrenic nerve-evoked endplate potentials (EPPs) were measured in P7 diaphragm muscle from *P2ry1* WT and mutant mice in response to basal (left panel) and high-frequency stimulation (HFS; middle and right panels). There was no difference in the amplitudes (24 ± 2.6 vs. 26.7 ± 1.5 mV; p=0.06; WT vs. mutant; *n*=4; *c*=16) of basal EPPs or in the amplitudes of EPPs at the end of a period of HFS. ***B***, Phrenic nerve-evoked muscle action potentials (APs) were measured in P7 diaphragm muscle from *P2ry1* WT and mutant mice in response to basal (left panel) and HFS (middle and right panels). There was no difference in the amplitudes (67.7 ± 7.1 vs. 67.8 ± 7.1 mV; p=0.49), the rise to peak (1.16 ± 0.2 vs. 1.07 ± 0.2 ms; p=0.33), or the time to 50% decay (2.42 ± 0.53 vs. 2.16 ± 0.82 ms; p=0.43) of basal APs, or in the percentage of successfully transmitted muscle APs at the end of a period of HFS (note the three successful APs in the WT and 2 APs in the mutant, at the end of HFS; arrowheads). ***C***, As shown in right panel of A, ending EPP heights are similar between genotypes (34 ± 8.6 vs. 34.3 ± 16.6 % initial EPP; p=0.48; WT vs. mutant; *n*=4; *c*=16). ***D***, As shown in right panel of B, the time to 50% failure in response to HFS is similar between genotypes, in all subtypes of fatiguability (3.7 ± 1.6 vs. 2.5 ± 2.4s for quick fatiguability; p=0.12; 22 ± 15.1 vs. 24.7 ± 13.5s for intermediate fatiguability; p=0.34; 37.3 ± 9 vs. 37.2 ± 6.8s for slow fatiguability; p=0.48; WT vs. mutant; *n*=4; *c*=27). ***E***, Muscle APs from the beginning (left) or end of a train of HFS from *P2ry1* WT (black; +/+) or mutant (gray; -/-) mice. Muscle AP rise-to-peak was lengthened (2.0 ± 0.6 vs. 2.7 ± 0.7ms; WT vs. mutant, *p*<0.05) and amplitude was reduced (58.1 ± 3.23 vs. 53.7 ± 4.5mV; WT vs. mutant, *n*=4, *c*=13; *p*<0.05) at the end of a train of HFS.

In order to assess postsynaptic function, we took advantage of BHC, a drug which blocks contraction of skeletal muscle without affecting neurotransmission and thus allows the electrophysiological and optical evaluation of muscle APs (Heredia et al., 2016). We first assessed individual nerve-evoked muscle APs and observed no differences between *P2ry1* WT and mutant mice (**Figure 4B**). In order to determine the effects of HFS on muscle APs, we initially examined neural transmission failure, or the failure to transmit a successful EPP into a muscle AP, by identifying the time at which less than half of the nerve stimuli were transduced into successful muscle APs. Similar to the adult diaphragm (Heredia et al., 2016), P7 diaphragm exhibited multiple muscle AP profiles in response to HFS, characterized by the occurrence of failed or subthreshold APs at different timepoints after stimulation, likely reflecting differential fatiguability. We were unable to detect differences in the time to neural transmission failure in any subtype (**Figure 4D**), consistent with the failure to detect differences in EPP or mEPP amplitude, which reflect the presynaptic release of and the postsynaptic response to ACh, respectively. However, when we examined the features of *successfully* transmitted muscle APs at different stages of HFS, we found that the amplitudes were shorter and durations longer in *P2ry1* mutant relative to WT mice (**Figure 4E**). These results suggest that the transduction of the muscle AP, rather than the transmission of the nerve impulse to the muscle, is affected in the absence of activity-induced, P2Y_1_R-mediated Ca^2+^ responses in TPSCs.

### Activity-induced, P2Y_1_R-mediated Ca^2+^ responses in TPSCs are required for the maintenance of muscle force in response to HFS

Because previous studies reported that impaired muscle APs are correlated with muscle fatigue (Juel, 1988), we evaluated muscle force in the P7 diaphragm of *P2ry1* mutant and WT mice. We used an optical measure of fiber shortening in whole diaphragm to measure muscle peak force and muscle fatigue (Heredia et al., 2016). When we examined the effect of 45s of 40Hz phrenic nerve stimulation, we detected no difference in the magnitude of peak contraction (**Figure 5A,B**). However, peak contraction was maintained for longer durations in *P2ry1* WT than mutant mice (**Figure 5B**). These results were also obtained in WT mice treated with the P2Y_1_R antagonist MRS2500, demonstrating that the acute inactivation of P2Y_1_R function is sufficient to enhance fatigue (**Figure 5C**). When we subjected the diaphragm to multiple bouts of HFS, each separated by a recovery period of 15 minutes, we found that the initial peak contraction, as well as ability to maintain peak contraction, were significantly reduced in *P2ry1* mutants (**Figure 5D**). Collectively, these data demonstrate that muscle fibers of the P7 diaphragm are more sensitive to fatigue induced by HFS in *P2ry1* mutant than WT mice. We next examined muscle fiber subtype in the diaphragm, since muscle AP failure profiles with different fatiguability were observed in response to HFS, and since the development of these subtypes reflects the endogenous pattern of nerve stimulation. Although earlier studies indicate that the development of these fiber subtypes first occurs at around P25 in rodent diaphragm (Zhan et al., 1998), we observed that both *P2ry1* WT and mutant mice contained all 4 basic fiber subtypes at P7, as assessed by immunostaining with myosin heavy chain (MHC) antibodies that selectively recognize each fiber subtype (Bloemberg and Quadrilatero, 2012). However, the relative percentage of each of these subtypes was indistinguishable between genotypes, suggesting that the enhanced fatigue in *P2ry1* mutants is not caused by a relative increase in fast-fatiguing subtypes of muscle fibers (**Figure 5E**).

**Figure 5.**
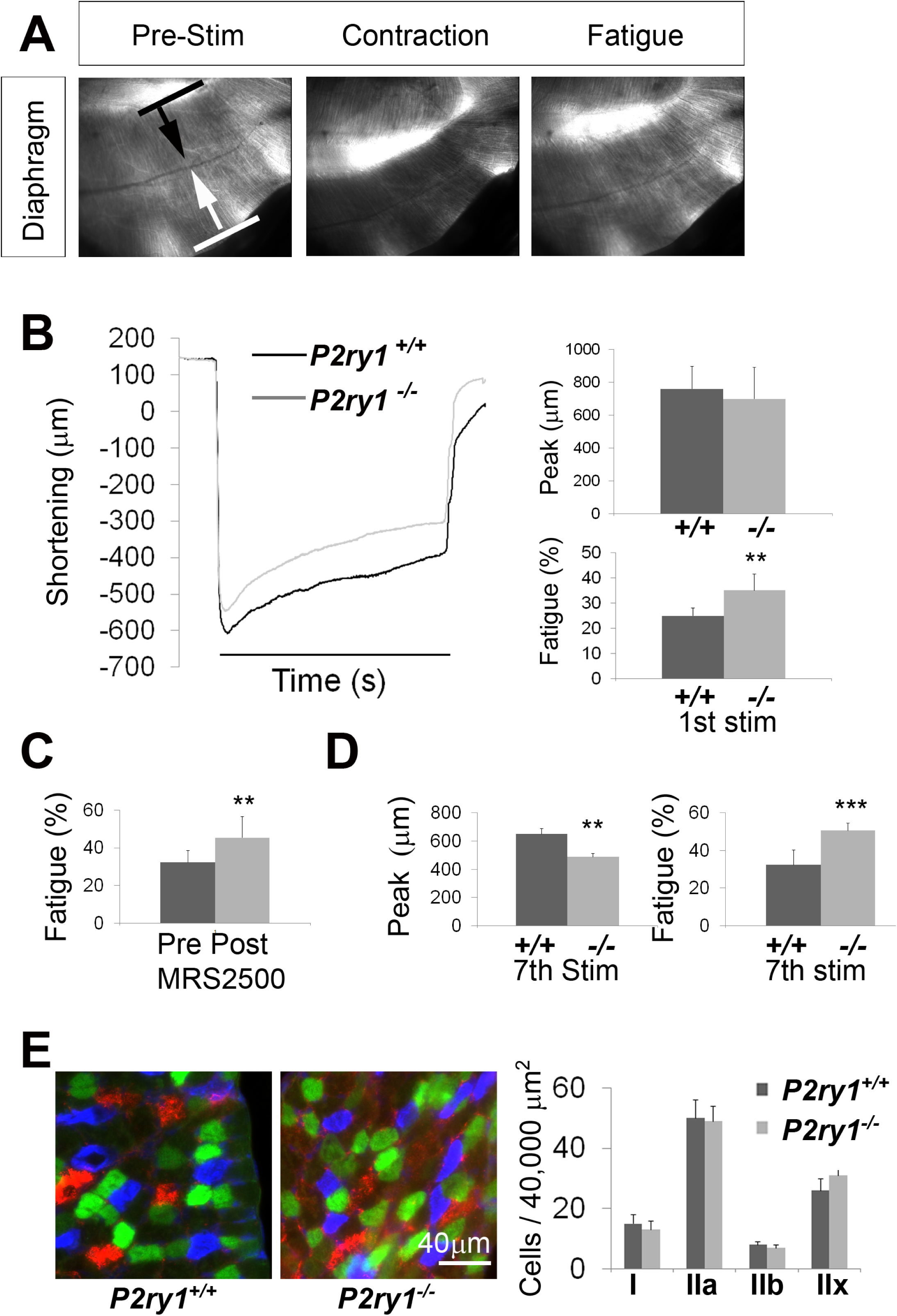
The loss of activity-induced Ca^2+^ responses in perisynaptic glia leads to enhanced muscle fatigue. ***A***, Images of a P7 hemi-diaphragm before phrenic nerve stimulation, at peak contraction, and during fatigue. Black and white arrows indicate sites used to compare fiber length changes. ***B***, The shortening of muscle fibers, as measured by the change in distance between the two sites indicated by arrows in A, is represented as a negative number. Peak length changes are similar between genotypes (760 ± 136 μm vs. 699 + 191 μm; p=0.22), but are maintained significantly less over time in *P2ry1* mutants (75 ± 3.1 vs. 66.9 ± 6.4% peak length change; *n*=9; *P2ry1* WT vs. mutant; *p*<0.01). ***C***, Fatigue is also enhanced by acute blockade of TPSC Ca^2+^ responses (73 ± 8.4 vs. 62.3 ± 8.3% peak length change; *n*=5; *P2ry1* WT vs. mutant; *p*<0.01). ***D***, Peak length changes in response to the seventh bout of HFS are reduced in *P2ry1* mutants (650 ± 38 μm vs. 487 ± 24 μm; *p*<0.01), as is fatigue (ending length change = 67.7 ± 8.1 vs. 49.5 ± 4% peak length change; *n*=5; *P2ry1* WT vs. mutant; *p*<0.03). ***E***, Image of transverse section of P7 diaphragm from *P2ry1* WT and mutant mice, stained with antibodies against myosin heavy chain (MHC) I (blue), MHC IIA (green) and MHC IIB (red) antibodies (left panel). No difference between genotypes was observed in the number of each MHC muscle fiber subtype.

A recent study demonstrated that neonatal TPSCs respond to distinct levels of nerve activity during the period of polyneuronal synapse elimination by modulating the magnitude of their Ca^2+^ response (Darabid et al., 2013). Together with the finding that TPSC processes separate competing nerve terminals from each other and from the postsynaptic muscle fiber during this period (Smith et al., 2013), these data suggest that TPSC Ca^2+^ responses may regulate this phenomenon. Further support for this idea comes from the finding that synapse elimination in the CNS is impaired in *P2ry1* mutant mice as well as in mice lacking activity-induced Ca^2+^ responses in astrocytes (Yang et al., 2016). In order to evaluate this hypothesis, we confirmed that Ca^2+^ responses were eliminated in the absence of P2Y_1_R signaling at P15, the age at which synapse elimination is largely complete. Similar to those at P7, TPSC Ca^2+^ responses at P15 were completely dependent on P2Y_1_R signaling (**Supplemental Figure 3A**). Fatigue was similarly enhanced, using both optical and tension measurements (**Supplemental Figure 3B**). However, when we examined NMJs by neurofilament immunohistochemistry to detect the numbers of innervating axons at individual NMJs at several ages between P7 and P15, we were unable to observe any differences (**Supplemental Figure 3C,D**). Together, these results suggest that activity-induced Ca^2+^ responses in TPSCs are not required for polyneuronal synapse elimination in the developing diaphragm.

### Muscle fatigue is more severely enhanced by high levels of potassium in *P2ry1* mutants lacking activity-induced, P2Y_1_R-mediated Ca^2+^ responses in TPSCs

A variety of mechanisms underlie muscle fatigue. In order to assess which of these might be affected by TPSC Ca^2+^ accumulation, we initially examined intracellular Ca^2+^ release within muscle cells. In order to investigate activity-induced Ca^2+^ signaling in whole populations of diaphragm muscle cells, we crossed Myf5-Cre to conditional GCaMP3 mice. In unparalyzed muscle under epifluorescence, we measured both fiber length changes and Ca^2+^ fluorescence intensities in response to HFS. Similar to the results obtained with brightfield recordings obtained above, MRS2500 enhanced the fatigue of a second bout of HFS relative to a first, compared to no treatment (data not shown). Peak intensities, time to 50% decay, and ending relative to peak intensities of Ca^2+^ transients were all significantly affected in response to HFS in the presence of MRS2500 (**Supplemental Figure 4**), suggesting that events upstream or concurrent with Ca^2+^ release mediate muscle fatigue caused by the absence of TPSC Ca^2+^ signaling.

Direct electrical stimulation of skeletal muscle produces fatigue, similar to indirect, nerve-mediated excitation. Interestingly, in response to high levels of extracellular potassium or high [K^+^]_o_, this activity-induced fatigue is enhanced, and muscle APs exhibit lower amplitudes and longer durations (Cairns et al, 2015), similar to the response of *P2ry1* mutants to nerve activity shown above. These results raise the possibility that activity-induced Ca^2+^ signaling in TPSCs protects against muscle fatigue by regulating K^+^ uptake by these cells and therefore persiynaptic [K^+^ _o_]. Interestingly, in a pair of studies on astrocytes in the hippocampus and Bergmann glia in the cerebellum, [K^+^]_o_ and K^+^ uptake were similarly affected by manipulations to activity-induced Ca^2+^ signaling (Wang et al., 2012a; 2012b). In order to test this idea, we first challenged diaphragms to [K^+^]_o_ greater or less than normal levels (5mM). These challenge experiments were modeled on those used to characterize the effects of hypo- and hyperkalemia on skeletal muscle function (Wu et al., 2011). We found that HFS-induced fatigue was ameliorated in response to low [K^+^]_o_ in *P2ry1* mutants (ending length change, 2.5mM [K^+^]_o_ vs. 5mM [K^+^]_o_ = 79.3 ± 13 vs. 68.3 ± 11.2% peak length change; *n*=3; *P2ry1* WT vs. mutant mice, *p*<0.01). Next, we performed the same experiments in response to 10mM [K^+^]_o_. Fatigue was disproportionately enhanced in *P2ry1* mutant, relative to WT mice (**Figure 6A**). The enhanced fatigue in these mutants was further revealed by multiple bouts of HFS; *P2ry1* mutant diaphragm was almost completely unable to contract after the second period of HFS in high [K^+^]_o_, in marked contrast to *P2ry1* WT diaphragm (**Supplemental Video 5; Figure 6A**). We examined whether this effect of high [K^+^]_o_ was caused by depolarization of the postsynaptic muscle membrane. After stimulation with several bouts of HFS in 5mM [K^+^]_o_, muscle cells were impaled and recorded before and after changing the [K^+^]_o_ to 10mM. While this caused a mild depolarization of the RMP in *P2ry1* WT mice, this effect was enhanced in *P2ry1* mutants (**Figure 6B**). Collectively, these results demonstrate that P7 diaphragm muscle cells lacking activity-induced, P2Y_1_R-mediated Ca^2+^ signaling in TPSCs are more sensitive to high [K^+^]_o_, suggesting that Ca^2+^ signaling modulates the response to [K^+^]_o_ in these perisynaptic glia.

**Figure 6.**
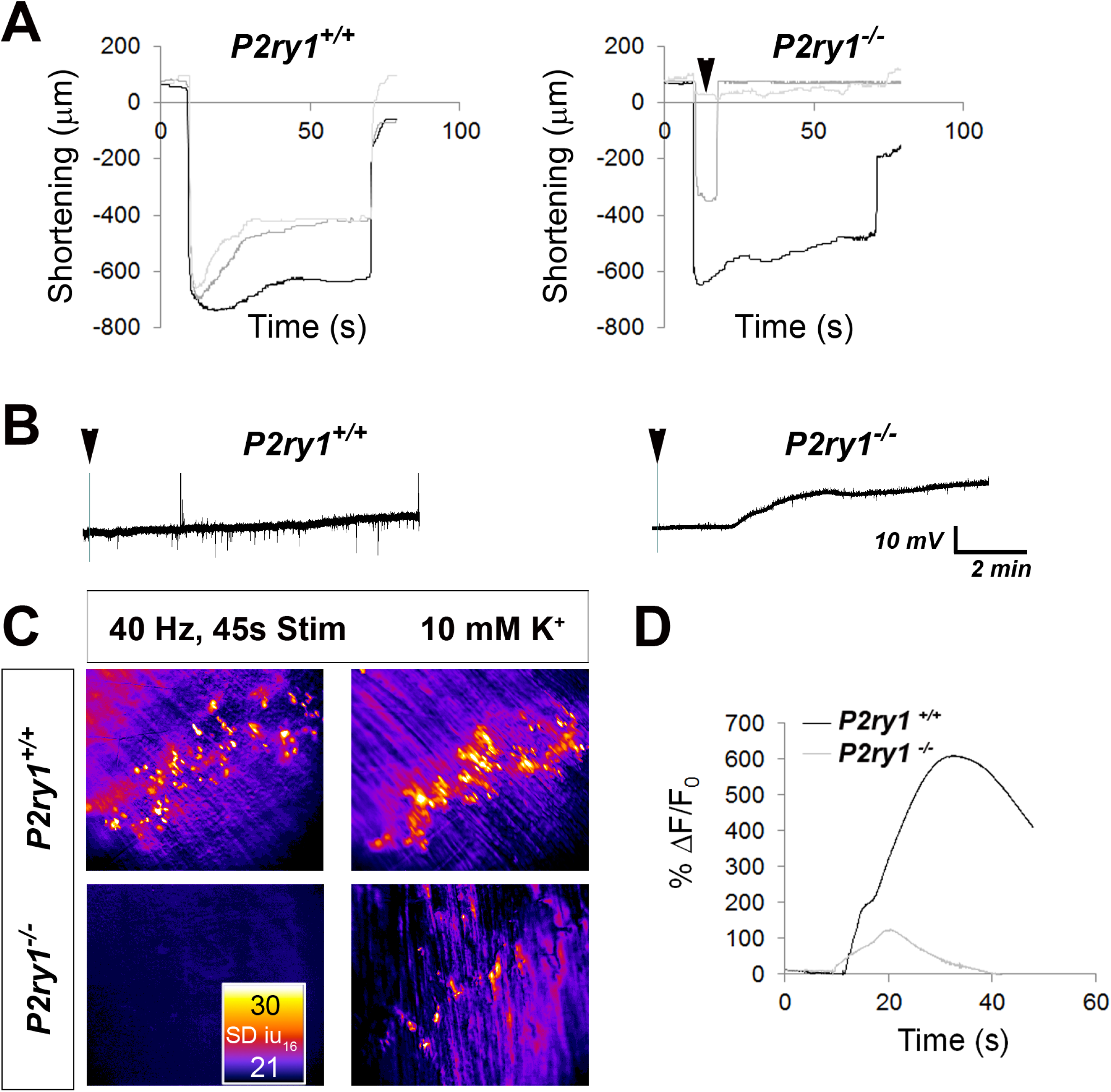
High [K^+^_o_] triggers greater muscle fatigue, greater muscle membrane depolarization, and reduced Ca^2+^ responses in the absence of activity-induced Ca^2+^ responses in perisynaptic glia. ***A,*** Muscle length changes from P7 diaphragm of *P2ry1* WT (left) and mutant (right) mice were first recorded in response to HFS in normal or 5mM [K^+^]_o_ (black traces), and then recorded in response to 2 bouts of HFS in 10 mM [K^+^]_o_ (dark and light grey lines = 1^st^ and 2^nd^ HFS, respectively). Peak length changes and fatigue are dramatically affected in *P2ry1* mutants by high [K^+^ o]: peak length in response to first HFS in 10mM [K^+^]_o_ as a percentage of peak length in response to control HFS in 5 mM [K^+^]_o_ = 96.3 ± 8 vs. 55.5 ± 13%; *n*=3; *P2ry1* WT vs. mutant mice, *p*<0.001; peak length change in response to second HFS in 10mM [K^+^]_o_ = 90.6 ± 11 vs.6.4 ± 5% control HFS; *n*=3; *P2ry1* WT vs. mutant mice; p<0.0005; asterisk indicates this almost complete failure to contract; ending length change of first HFS in high 10mM [K^+^]_o_ = 62.3 ± 8 vs. 0% peak length change; *n*=3; *P2ry1* WT vs. mutant mice, *p*<0.0001). ***B***, Effect of high [K^+^]_o_ on resting membrane potential (RMP). Representative muscle cell recording before and after (arrowhead) [K^+^]_o_ was changed from 5-10mM in P7 diaphragm from *P2ry1* WT (left) and mutant (right). ***C***, SD intensity maps of TPSCs in response to HFS (left panels) in *P2ry1* WT (upper) and mutant (lower) mice and in response to subsequent treatment with 10mM [K^+^]_o_ (right panels). Markedly fewer TPSCs responded to elevated [K^+^]_o_ in *P2ry1* mutants. ***D***, Ca^2+^ transient intensities of TPSCs responding to 10mM [K^+^]_o_ were significantly reduced in *P2ry1* mutants (30 ± 7.3 vs. 22.9 ± 4 dB WT vs. mutant, *n*=3; *p*<0.001); fire CLUT heatmap in SD iu_16_.

### P2ry1 mutants lacking activity-induced, P2Y_1_R-mediated Ca^2+^ responses in TPSCs exhibit a reduced potassium-induced Ca^2+^ response

To test if TPSCs take up K^+^ at the NMJ, we took advantage of an observation that raising [K^+^]_o_ to 10mM resulted in a robust TPSC Ca^2+^ response. This was not mediated by indirect depolarizing effects of [K^+^]_o_ on the phrenic nerve, leading to an activity-induced, P2Y_1_R-mediated Ca^2+^ response, because it was still observed in the presence of doses of tetrodotoxin that blocked neurotransmission, and because it was also observed in *P2ry1* mutants completely lacking activity-induced Ca^2+^ responses. Interestingly, Ca^2+^ responses induced by high [K^+^]_o_ were not caused by depolarization of TPSCs, because they were still observed when external Ca^2+^ was removed, but rather by release from intracellular stores, because they were abrogated after treatment with CPA (n=3; data not shown).

If K^+^ uptake is affected in TPSCs of *P2ry1* mutants lacking activity-induced Ca^2+^ signaling, then this [K^+^]_o_-induced Ca^2+^ response in these cells should in turn be reduced. In order to test this idea, we subjected diaphragms to several bouts of HFS, separated by 15 minutes, to mimic the effects on muscle fatigue described above, then assessed the effects of high [K^+^]_o_ on Ca^2+^ signaling. In contrast to *P2ry1* WT mice, mutants showed a markedly reduced Ca^2+^ response to 10mM [K^+^]_o_ (**Figure 6C,D**), suggesting that K^+^ uptake is indeed impaired in *P2ry1* mutants. This failure of TPSCs to buffer perisynaptic [K^+^]_o_ may contribute to the enhanced muscle fatigue that occurs in *P2ry1* mutants lacking activity-induced TPSC Ca^2+^ responses. Additionally, these results suggest that TPSC Ca^2+^ responses, and consequently K^+^ uptake, are positively regulated by nerve activity through feedforward (i.e., neurotransmitter-mediated stimulation) and feedback mechanisms (i.e., by suprathreshold [K^+^]_o_ itself)

## Discussion

Our results using Wnt1-GCaMP3 mice demonstrate that activity-induced Ca^2+^ signaling in neonatal TPSCs of the diaphragm is mediated by P2Y_1_R activation by nerve-derived adenine nucleotides. The absence of Ca^2+^ signaling within TPSCs does not appear to affect the structural and molecular development of the NMJ, nor does it alter the presynaptic release of neurotransmitter, but rather affects the postsynaptic AP during sustained HFS, where longer, smaller APs were correlated with a failure to maintain peak muscle force. Because previous studies observed that administration of high K^+^ induced similar effects on muscle APs subjected to fatiguing muscle stimulation (Cairns et al., 2015), we examined muscle fatigue in response to this treatment and found that it was enhanced to a greater degree in *P2ry1* mutants lacking activity-induced Ca^2+^ signaling. This heightened susceptibility to high [K^+^]_o_ may be caused by impaired K^+^ uptake by *P2ry1* mutant TPSCs, as these cells exhibited a markedly reduced release of Ca^2+^ from intracellular stores in response to high [K^+^]_o_ treatment. Collectively, these results suggest that activity-induced, P2Y_1_R-mediated Ca^2+^ signaling in TPSCs influences muscle fatigue by regulating perisynaptic [K^+^]_o_.

The current study represents the first evaluation of Ca^2+^ responses using genetically encoded calcium indicators in perisynaptic glia at the NMJ. The onset after nerve stimulation and time to peak intensity of Ca^2+-^ transients using this method are similar to those of published studies using Ca^2+-^ sensitive dyes in TPSCs loaded with Fluo-3 as well as astrocytes expressing GCaMP3 (Jaromi et al., 1992; Reist and Smith, 1992; Darabid et al., 2013; Akerboom et al., 2013), demonstrating that this genetic technique is a valid tool to measure these responses in large populations of TPSCs. The most striking finding from this study is the complete dependence of TPSC Ca^2+-^ responses on a single GPCR, P2Y_1_R. Therefore, despite the fact that bath administration of a multiple substances induces widespread Ca^2+^ signaling in neonatal TPSCs, activity-induced responses are only mediated by adenine nucleotides. In contrast to these results, studies of adult TPSCs support a role for ACh and other factors in mediating these responses (Darabid et al., 2014). Therefore, the early exclusive dependence on P2Y_1_R-activating adenine nucleotides may broaden over time. Alternatively, TPSCs at the NMJs of the diaphragm may continue to depend exclusively on P2Y_1_R signaling. At the oldest ages at which we were able to examine population responses (P15-P20), activity-induced Ca^2+^ signals were completely dependent on the P2Y_1_R pathway, supporting this latter idea. Moreover, the prevention of ACh diffusion to perisynaptic mAChRs by cholinesterase is unlikely to represent a developmentally transient response. Indeed, a recent report described a functional role for TPSC-derived BChE at the adult NMJ (Petrov et al., 2013).

Our results fail to support the idea that TPSC Ca^2+^ responses affect the presynaptic release of ACh, in contrast to previous studies (Robitaille, 1998; Castonguay and Robitaille, 2001). NMJs at the diaphragm do exhibit dynamic changes of ACh release such as facilitation and depression (Vautrin et al., 1993), arguing against the absence of plasticity at diaphragm NMJs as an explanation of these differences. Rather, they may be attributable to different species, muscle, age or technique. Interestingly, using a similar genetic approach to study the effect of eliminating activity-induced Ca^2+^ responses in astrocytes neurotransmitter release was unaffected (Agulhon et al., 2010). On the other hand, postsynaptic function was affected at the NMJ in response to HFS in *P2ry1* mutants. Whereas the number of successful muscle APs, reflecting the presynaptic release of and response to ACh, was not different in response to HFS, the characteristics of these APs changed significantly in response to prolonged nerve stimulation. These results suggest that muscle APs may not be transduced as efficiently in *P2ry1* mutant mice. The reduced intensity of muscle Ca^2+^ transients in response to pharmacological blockade of P2Y_1_Rs supports this contention, as does the enhancement of muscle fatigue in response to pharmacological or genetic inhibition of this pathway. These effects may result from the absence of P2Y_1_R function in muscle, as the mice used in this study were constitutive mutants. For example, Choi et al. (2001) found that stimulation of chick muscle fibers with 100μM ATP triggered Ca^2+^ release from intracellular stores, a response blocked by the pan-P2 blocker suramin. Together with the expression of P2Y_1_R by muscle, these findings suggest that nerve-derived ATP may modulate intracellular muscle Ca^2+^ levels and thus muscle fatigue. However, we failed to observe Ca^2+^ release in response to ATP in the muscle cells of Myf5-GCaMP3 mice (data not shown). Moreover, Ca^2+^ signals in TPSCs were observed in response to treatment with lower doses of ADP/ATP (10-20 μM) than those used to evoke these signals in chick muscle cells. Finally, Ca^2+^ signals in TPSCs were blocked after P2Y_1_R was blocked by specific pharmacological and genetic tools, rather than the pan-P2R blocker suramin (Choi et al., 2001). Thus, we favor the idea that the reduction of muscle cell Ca^2+^ mobilization in *P2ry1* mutants is caused by the absence of this protein in TPSCs rather than in muscle cells.

The effects of HFS on muscle APs suggested the possibility that perisynaptic [K^+^]_o_ was dysregulated in *P2ry1* mutants. Supporting this idea, treatment with high [K^+^]_o_ enhanced muscle fatigue to a greater extent in *P2ry1* mutant than in WT mice. Together with a report that intense exercise increases [K^+^]_o_ in muscle to 10-14 mM (Mohr et al., 2004), a level sufficient to cause muscle fatigue (Sjøgaard et al., 1990; but see Shushakov et al., 2007), these data indicate that muscle fatigue is enhanced in *P2ry1* mutants as a result of elevated perisynaptic [K^+^]_o_, caused by a failure of TPSCs to spatially buffer or take up K^+^ (Kofuji and Newman, 2004). In order to test this idea, we examined the effects of treatment with high [K^+^]_o_, reasoning that if K^+^ uptake mechanisms in TPSCs were impaired in *P2ry1* mutants lacking activity-induced Ca^2+^ signaling, these effects would be diminished. Indeed, we found that Ca^2+^ responses to 10mM [K^+^]_o_ were markedly reduced in these mutants, providing indirect evidence that K^+^ uptake was impaired. Interestingly, elevation of [K^+^]_o_ to 20 mM also induced a robust increase of intracellular Ca^2+^ in cultured astrocytes (Duffy and MacVicar, 1994). However, in contrast to the current study, this response was mediated entirely by external influx through voltage-gated Ca^2+^ channels (VGCC). Therefore, K^+^-induced Ca^2+^ responses in TPSCs do not appear to result from depolarization-mediated ingress through VGCC, but rather by the release from intracellular stores. Together, these data suggest that activity stimulates perisynaptic K^+^ uptake in TPSCs by both feedforward Ca^2+^ responses initiated by neurotransmitter and feedback signals initiated by high [K^+^]_o_ itself.

The intracellular uptake of K^+^ against its concentration gradient has been demonstrated in Müller glia in the retina (Newman et al., 1984) and is mediated by several mechanisms, including inwardly-rectifying potassium channels (Kir), Na^+^, K^+^ ATPases, and Na^+^, K^+^ Cl^-^ cotransporters. The importance of K^+^ uptake has been demonstrated by several genetic studies. For example, mice lacking Kir4.1 in astrocytes exhibit impaired K^+^ and neurotransmitter uptake, leading to seizures, ataxia and early lethality (Djukic et al., 2008). Na^+^, K^+^ ATPase activity in astrocytes has also been implicated in regulating perisynaptic [K^+^]_o_. For example, Wang et al. (2012) observed that activity-induced Ca^2+^ responses in astrocytes are required for K^+^ uptake through an ouabain-sensitive Na^+^, K^+^ ATPase activity (Wang et al., 2012a). Future studies will determine which if any inward K^+^ conductance is expressed in and stimulated by activity within TPSCs, as well as the mechanisms by which increases of intracellular Ca^2+^ lead to enhanced K^+^ uptake. Of note, it has been established that nonmyelinating Schwann cells in sympathetic nerves possess Kir currents that are sensitive to neural activity (Konishi, 1994).

In addition to [K^+^]_o_, the extracellular concentrations of other ions are dysregulated during muscle fatigue, including [Na^+^]_o_ and [H^+^]_o_ (Allen et al., 2008). Because astrocytes express an abundance of transporters and ion channels that modulate the levels of these ions in response to neuronal activity, TPSCs may similarly regulate these ions in response to stimulation of muscle, in addition to [K^+^]_o_. On the other hand, muscle itself is equipped for this role, expressing an abundance of ion channels and transporters along the extensive t-tubule system that regulate muscle membrane potential and excitability during activity (Fraser et al., 2011). However, the importance of ionic homeostasis at the NMJ may depend on additional mechanisms, such as those proposed here. For example, the sensitivity of Na_v_1.4, which is localized in a restricted region in the depths of the postsynaptic junctional folds, to the effects of perisynaptic [K^+^]_o_, may require enriched expression of [K^+^]_o_ buffering proteins by TPSCs.

In summary, we have utilized the diaphragm of neonatal *P2ry1* mutant mice as a model to explore the functional significance of activity-induced Ca^2+^ signals in perisynaptic glia. We found that in the absence of purinergic signaling, postsynaptic rather than presynaptic function was altered, leading to enhanced muscle fatigue in response to HFS. These effects were correlated with elevated [K^+^]_o_ and reduced responsivity to [K^+^]_o_, suggesting that activity-induced Ca^2+^ responses in TPSCs regulate perisynaptic [K^+^]_o._ Future studies will determine the mechanisms underlying K^+^ uptake and [K^+^]_o_-mediated Ca^2+^ accumulation in TPSCs. Such mechanisms may represent important translational targets in diseases with altered [K^+^]_o_. For example, in patients with hyperkalemic periodic paralysis, a genetic disorder caused by *Scna4* mutations and characterized by elevated [K^+^]_o_, stimulation of Ca^2+^ signaling and subsequently [K^+^]_o_ buffering within TPSCs may enhance neuromuscular function.

## Materials and Methods

### Ethical Approval and Use of Mice

*P2ry1* mutant, GCaMP3 or GCaMP6f conditional knockin, and *Wnt1-Cre* and *Myf5-Cre* transgenic mice were all purchased from Jax. *P2ry1* null mutant mice were backcrossed into the C57/Bl6 strain several times before crossing to other strains, each of which is maintained in the C57/Bl6 strain. We could find no difference in any experiment between *P2ry1*^*+/+*^ and *P2ry1*^*+/-*^ mice, so we pooled these samples and denoted them all in the text as “*WT*.” Similarly, we found no difference between male and female *P2ry1*^*+/-*^ mice, so we pooled these samples. In order to generate *P2ry1* mutants expressing GCaMP3 in Schwann cells, we generated *Wnt1-Cre*; *P2ry1*^*+/-*^ and *Rosa26-GCaMP3*^*flox/flox*^; *P2ry1*^*+/-*^ mice and crossed them, such that all *P2ry1* mutant and heterozygote mice expressed only one copy each of Cre and GCaMP3. We used a slightly modified common 3’ WT primer to genotype *P2ry1* mutant mice (ATT TTT AGA CTC ACG ACT TTC) and the recommended primers by Jax for all other alleles. All studies were performed with animals aged postnatal day 7 and 15 (P7, P15). To verify knockouts, we performed RT-PCR on muscle-derived RNA with primers against a 300-bp fragment of *P2ry1* (5’: CTG TGT CTT ATA TCC CTT TCC, 3’: CTC CAT TCT GCT TGA ACT C). Animal husbandry and experiments were performed in accordance with the National Institutes of Health *Guide for the Care and Use of Laboratory Animals* and the IACUC at the University of Nevada.

### Drugs

The following reagents were used at the following concentrations: P2Y_1_R agonists ATP and ADP (Sigma; 10 or 20μM); P2Y_1_R antagonist MRS2500 (Tocris; 1μm); P1R agonist adenosine (Sigma; 100μM); pan-P2 antagonist suramin (Sigma; 100μM); pan-muscarinic agonist muscarine (Sigma; 10μM); pan-muscarinic blocker atropine (Sigma; 10μM); pan-nicotinic agonist nicotine (Sigma; 50μM); pan-nicotinic antagonist curare (Sigma; 200μM); pan-cholinesterase inhibitor neostigmine (Sigma; 1μM); sarco-/endoplasmic reticulum Ca^2+-^ATPase (SERCA) inhibitor, cyclopiazonic acid (Sigma; CPA; 10μM); potassium chloride (Sigma; 2-10 mM); GIIIb μ-conotoxin (Peptides International; 2.3μM); skeletal muscle myosin-blocker BHC (Hit2lead; 100μM); 488-, 594-, 633-conjugated-α-bungarotoxin (α-BTX; Biotium; 1μg/mL).

### Calcium Imaging

The diaphragm of Wnt1-GCaMP3 mice was illuminated with a Spectra X light engine (Lumencor). In order to quantify maximal fluorescence (F_max_) exhibited by GCaMP3 in Schwann cells, 30μM CPA was added to deplete sarcoplasmic reticular Ca^2+^ stores (Heredia et al., 2016). Image sequences were captured using an Andor Neo sCMOS camera and a Windows-based PC using Nikon NIS Elements 4.1. Image sequences were recorded at 25 frames per second, and were exported as 8-bit TIFF files into custom-written software (Volumetry G8d). A Gauss filter (3x3 pixel, sd = 1.0) was applied to reduce camera noise, and motion-correction routines were used to stabilize neural and Schwann cell elements in the movie (see Hennig et al, 2015). Changes in background fluorescence were stabilized by subtracting the average intensity near the main phrenic nerve branch. An average intensity image was generated before stimulus application (“Pre-stim” ∼ 1s) to quantify basal Ca^2+^ levels in Schwann cells. These images are presented using a blue->green color lookup table (CLUT). This image was subtracted from the entire movie, thereby filtering out static fluorescent structures and displaying only objects that changed their intensity, i.e., Ca^2+^ transients. A number of statistical maps (stat maps) were used to portray and analyze the pattern of activity-induced Ca^2+^ transients in TPSCs. The main stat map type used to portray the amplitude of Ca^2+^ transients in TPSCs was the standard deviation (SD) map. This map was calculated in similar fashion to the average intensity image, except the standard deviation of 16-bit fluorescence intensity units (SD iu_16_) at every pixel prior to the application of the stimulus (0.5-1.0s) extending to 60s was calculated. SD maps were color coded using a “Fire” CLUT. Traces of fluorescence intensity were generated from movies and presented as changes in fluorescence with respect to initial fluorescence (ΔF/F_avg(prestim)_ or ΔF/F_o_). Peak intensities of Ca^2+^ transients were calculated as a signal-to-noise ratio (SNR) in which peak standard deviation values were divided by the prestim standard deviation value. The log_10_ of this ratio was generated and multiplied by 20 to standardize the SNR as decibels (dB). Drugs were either bath applied in proximity to the motor endplate or pressure injected (PDES-O2DX; NPI Electronic). Drugs dissolved in DMSO were either perfused in or diluted in 1 mL of Kreb’s-Ringer’s before bath application, as bath application of small volumes of DMSO (∼8μl into 8ml chamber) caused fluorescence within Wnt1-GCaMP3-expressing TPSCs. For experiments with altered potassium, a stock solution of 3M [KCl] was added to the bath to change the concentration from 5mM to 10mM immediately prior to imaging.

### Electrophysiology

Diaphragms were dissected and pinned on a 6-cm Sylgard-coated dish containing oxygenated Krebs-Ringer’s solution at RT according to standard procedure. Stimulation and recording of intracellular potentials were performed as described (Heredia et al., 2016). Briefly muscle APs were recorded after treatment with 100μM BHC for 30 minutes, followed by 30 minutes of washing. Endplate potentials (EPPs) were recorded after treatment with the Na_v_1.4 antagonist μ-conotoxin GIIIb (μ-CTX; 2.3μM). Signals were amplified, digitized, recorded and analyzed as described (Heredia et al., 2016). Only muscle fibers with resting membrane potentials between -60 and -75mV were included for analysis. Stimulation episodes of the phrenic nerve over 10Hz were separated by 30-minute rest periods to allow recovery. In order to calculate percent failure (APs) or percent transmitter release rundown (EPPs), the average of three potentials at a particular timepoint (e.g., the time at which fewer than half of nerve stimuli produced a successful muscle AP) was taken and expressed as a percent of the average of the first three potentials. For experiments with altered KCl, NaCl was adjusted accordingly to maintain the same Cl^-^ concentration before being perfused into the dish.

### Fatigue

Tension recording of muscle force in diaphragm strips (P15) or video recording of muscle shortening in hemidiaphragms (P7, P15) was performed as described (Heredia et al., 2016). For experiments with altered KCl, NaCl was adjusted accordingly to maintain the same Cl^-^ concentration before being perfused in.

### Immunohistochemistry

Antibodies against GFP (Rockland), S100 (Dako), synaptophysin (Santa Cruz), neurofilament (Millipore) and acetylcholinesterase (kindly provided by P. Taylor, UCSD) were used at 1/1000 in PBS containing 1% triton-X and 10% fetal bovine serum to detect proteins in fixed, whole-mount diaphragms. Fluorescently-conjugated α-BTX and fasciculin-2 were added with secondary antibodies. Tissues were confocally imaged with an Olympus Fluoview 1000. For myosin heavy chain staining, muscles were fresh-frozen, cut at 16μm, and immediately incubated without fixation in PBS with primary antibodies as described (Heredia et al., 2016).

### Electron Microscopy

P7 mice were transcardially perfused in 1.5% glutaraldehyde, 2% paraformaldehyde in 0.1M sodium cacodylate. The costal diaphragm was dissected and incubated in fixative at 4^0^C overnight and then in rinse for several hours at 4^0^C. Samples were post-fixed in 2% osmium tetroxide, dehydrated, incubated in propylene oxide, embedded in Spurr’s resin and polymerized at 60°C overnight. Ultrathin sections were cut at 90μm and stained with uranyl acetate followed by lead citrate. Sections were photographed or digitized using a Phillips CM10 transmission electron microscope equipped with a Gatan BioScan digital imaging system.

### Statistics

Power analyses were performed using G*power 3.010 to determine the numbers of *P2ry1* wild-type (WT) and mutant mice required. For example, to determine the number of mice (*n*) and cells (c) to analyze for electrophysiological recordings, a power of 0.8, significance or alpha of 0.05 and effect size or Pearson’s r of 3.6 was used. Thus, for these experiments, data was generated from *c*=3 per animal or more, and from *n*=3 or more. In this case, each *c* and each *n* are biological replicates. Differences between *P2ry1* mutant and WT mice were assessed by unpaired Student’s *t*-tests between two independent means, assuming equal variance. A *P* < 0.05 was considered significant.

## Acknowledgements

This work was supported with funds from the National Institutes of Health (NIH): GM103554 and GM110767 (T.W.G); National Center for Research Resources (5P20RR018751-09) and the National Institute of General Medical Sciences (8P20 GM103513-09).

## Contributions

D.J.H designed, carried out and interpreted the experiments in Figures 1-4; 6-10. C-Y F. carried out, interpreted and drafted the writing of the electron microscopy experiments in Figure 5. G.W.H. analyzed, interpreted, drafted and revised the writing of the experiments in Figures 2-3.T.W.G designed and interpreted the experiments in Figures 1-10. D.J.H. drafted and T.W.G. edited the manuscript.

All authors approve of the final version and agree to be held accountable for all aspects of the work.

## Statement of competing interests

The authors declare no competing interests.

Correspondence and requests for materials should be addressed to T.W. Gould

